# *Caenorhabditis elegans*: a model to study the anthelmintic effects of polyphenolic compounds on the fertility of parasitic gastrointestinal nematodes?

**DOI:** 10.1101/2020.09.09.289637

**Authors:** C Arroyo-Lopez

**Author notes:** Corresponding author: Celia, Arroyo-López, Ph.D.

## Abstract

We set up a Cae*norhabditis elegans* model to extrapolate anthelmintic (AH) effects of commercial polyphenols to related gastrointestinal zoonotic species. We compared the practical convenience of S liquid to solid agar Nematode grown medium in daily reproduction patterns and total brood size. Solid agar resulted a highly effective, reproducibly medium, in a less costly and rapid time manner. A minimum of three replications per monomer concentration are suggested to get a robust statistical analysis. Flavonoids Quercetin and Rutin potentially decrease brood sizes and percentage of development, with the exception of 20μM concentration. Flavanols resulted heterogeneous. In a dose-dependent manner Catechin 20μM significantly decreased egg output, and egg hatching in a 15% on respect to control values. Contrary, Epigallocatechin-gallate, seems to stimulate brood sizes and egg-hatching, however, EGCG10μM decreased reproduction but with no in a significative manner. We found a nematocidal effect on the reproductive parameters of *C. elegans* exposed to the chalcones Phloretin and Phloridzin, and Flavones Flavone and Genistein. A significant general decrease on reproductive parameters were found, particularly significant for Genistein 20μM in the percentage of hatching respect to control. Effects of time schedule and monomer concentration were found for the Hydrolysable tannins Gallic and Tannic acid. Tannic acid showed effectiveness against egg-hatching. The spectrum of percentages of change compared to each blank resulted negative decreasing fertility values, but for GA5, associated with a net increase of larvae hatched.

^1^ AH, GIN, CTS, HTs Q) Quercetin, (R) Rutin, (C) (+)-Catechin hydrate, (EGCG) (-)-Epigallocatechin Gallate, (PTN) Phloretin, (PDN) Phloridzin Dihydrate, (GE) Genistein, (FLA) Flavone, (GA) Gallic acid and (TA) Tannic acid., G1 (adult worm originally seeded per plate and monomer concentration at time 0. Two replications were performed per plate). Solvents: PBS (Phosphate Buffer Solution), CH4O (Methanol 0,002%) L3

## Introduction

Gastrointestinal nematodes (GINs) represent an important disease within livestock’s’ productive systems. The generalization and overuses of therapeutic drugs have prompted a high prevalence of multi-drug resistances within target parasitic population impacting small ruminant production systems (Torres-Acosta and Hoste, 2008), animal health (Jackson et al., 2012; Kaplan and Vidyashankar, 2012; Waller, 2007), persuading environmental risks (Beynon, 2012; Horvat et al., 2012; Wagil et al., 2015), and consumers concerns (Whelan et al., 2012). Resistances against the major families of anthelmintics represent a worldwide phenomenon to overcome to preserve the effectiveness of the anthelmintics (AHs). The development of environmentally friendly strategies based on the use of bioactive plants represent an alternative to direct or indirect impact the biology of parasites, and positively sway host welfare and consumers expectations (Durmic and Blache; Rochfort et al., 2008). The diversity of effectiveness and mode of action of traditional medicinal plants have also been studied *in vitro* and *in vivo* in the model *C. elegans* (Tang and Halliwell, 2010). Natural products as plant extracts have been traditionally used as a complement to traditional drugs or as the therapy itself (Fabricant and Farnsworth, 2001; Spiegler et al., 2017). Advances in alternative AHs research head for medicinal plants exploitation (Spiegler et al., 2016). Interest in locally grown plants rich in tannin content, and other potential secondary metabolites with AH properties (Vargas-Magaña et al., 2014) represent an ideal option for Green farming nowadays (Katiki et al., 2013). Last decade studies have demonstrated the AH effectiveness of tannin-rich resources on stages from egg to adulthood. However, GINs also showed different degrees of sensibility depending on the particular stage of their life cycle (Spiegler et al., 2017) and target species. A great number of publications screened the effectiveness of plant polyphenolic compounds against GINs (Athanasiadou et al., 2009; Hoste et al., 2006; Hoste et al., 2012; Hoste et al., 2015; Ketzis et al., 2006; Nguyen et al., 2005; Spiegler et al., 2017). For several years, our laboratory was involved in testing the AH efficacy of diverse Mediterranean tannin-rich plants (Manolaraki et al., 2010), Sainfoin (*Onobrychis viciifolia*), Carob (*Ceratonia siliqua*) (Arroyo-Lopez et al., 2014), or Tropical plants as Tzalam (*Lysiloma latisiliquum*) (Brunet et al., 2008b; Martínez-Ortíz-de-Montellano et al., 2013; Martínez-Ortíz-de-Montellano et al., 2010). Our trials were performed against the most common parasitic species within goats and sheep as *Haemonchus contortus* or *Trichostrongylus colubriformis* at different stages of their life cycle (see review (Hoste et al., 2015)). For example, *In vitro* studies indicated the effectiveness of *Onobrychis viciifolia* on egg-hatching and GIN larval migration inhibition (Barrau et al., 2005; Brunet et al., 2008b; Manolaraki et al., 2010). Furthermore, *in vitro* studies pointed towards the inhibition of pre-infective L3 exsheathment (Brunet and Hoste, 2006), and the establishment of infective L3 stage on gut mucosa (Brunet et al., 2008a). Additionally, *In vitro* and *In vivo* studies, showed effects on motility and survival rates of the adult population (Manolaraki et al., 2010). Aggregates of Tzalam and *O. viciifolia* extracts on adult *H. contortus* stages were observed *in vivo* and *in vitro* potentially, limiting their nutritional, reproductive functions, and motility (Martínez-Ortíz-de-Montellano et al., 2013). The anthelmintic challenges nowadays are to prevent resistances, to establish adequate dosages depending on pharmacodynamics, and the identification of the perfect timing for treatments. To reach this goal, the comprehension of the mechanism of action of the AHs on each stage, and the identification of potential molecules involved is essential. However, monitoring parasites and anthelmintics in the course of infection implies the killing and dissection of hosts as well as complications of keeping alive adults outside their host (Katiki et al., 2011; Sangster and Gill, 1999). *Caenorhabditis elegans* (Maupas, 1900) offers then multiple advantages as a translational model to evaluate AHs effects (Burns et al., 2015). Nevertheless, within the field of parasitology, *C. elegans* model remains controversial. Firstly, due to the natural variances on life cycles between parasitic and the free-living nematodes (Geary and Thompson, 2001), and secondly due to the different efficacy of the major types of the commercialized anthelmintic drugs against the most common parasites (Hu et al., 2013). Nevertheless, AHs exert similar nematocidal effects despite parasitic or free-living condition (Holden-Dye and Walker, 2014). *C. elegans* is an easy-breeding nematode, bacterivorous and hermaphrodite female (Wood, 1988). Its cell lineage, genes, anatomy, and developmental biology have been traced, providing a suitable model for genetics, biochemistry, physiology or neuroscience (Hiepe et al., 2006; Muschiol et al., 2009). It is also proposed as a good model for anthelmintic screening (Burns et al., 2015), based on the similarities on their physiology, pharmacology (Geary and Thompson, 2001; Holden-Dye and Walker, 2007), genomic organization, and the homology of some specific genes (Mitreva et al., 2005) with other members of the phyla Nematoda independently to their habitats. Besides, some authors remark a close association between the Order Rhabditida (*C. elegans*) with the Order Strongylida which encloses *H. contortus* and *T. colubriformis* responsible for the major trichostrongyliasis on ruminants (Geary and Thompson, 2001, Laing et al., 2013). Likewise, *C. elegans’* Dauer larvae shares analogies with the infective larvae (L3) from GINs, and similarities in the glycoproteic composition and structure of the cuticle with *H. contortus* (Bürglin et al., 1998; Fetterer, 1989), and ion channels from the nematode nervous system (Laing et al., 2013). Based on these analogies, anthelmintic effects observed against *C. elegans* might also be expected in related nematodes as *H. contortus* (Thompson et al., 1996). In a less costly and rapid manner, different compounds can be evaluated and extrapolate to related nematode parasitic species (Katiki et al., 2011; Thompson et al., 1996). Besides the nematocidal effect, effects on reproduction, behavior and/or locomotion can be examined (Rand and Johnson, 1995; Sangster and Gill, 1999; Strange, 2006). Until now, many other parasitologists have also included this model in the screening of plant extracts (McGaw et al., 2007; Waterman et al.), purified fractions, plant compounds (Katiki et al., 2011; Smith et al., 2009) and novel molecules tested in a liquid medium (Mathew et al., 2016). Based on published reports, the aim of this work was: 1) to set up an appropriate, easily reproducible, nematode grown medium and protocol for the evaluation of *C. elegans* fecundity rate. 2) to screen lethality and reproductive alterations on *C. elegans* after the exposure to polyphenolic compounds, 3) to evaluate AH effectiveness of some flavonoids, precursors of flavonoids and Hydrolysable Tannins (HTs).

## Ethics statement

The experiments performed with *C. elegans* followed the specific ethics required. No higher vertebrate animals were implied in these trials.

## Material and Methods

### Localization

*Escherichia coli* OP50 batches, monomer solutions, and media preparation requiring aseptic conditions were set up at the Unité Mixte de Microbiologie Moléculaire, Institut National de la Recherche Agronomique, Ecole Nationale Vétérinaire de Toulouse, (France). *Caenorhabditis elegans* culture, maintenance, and anthelmintic assays were performed at the UMR 1225 INRA/DGER Interaction Hôtes-agents Pathogènes, Institute National de la Recherche Agronomique, Ecole Nationale Vétérinaire de Toulouse, (France).

### Monomers and solvents

Phenolic compounds were purchased from Sigma-Aldrich (now MERK): (flavonoids) Flavonoids: Quercetin ≥95% (Q) (Q4951) and Rutin hydrate ≥94% (R) (R5143), Flavanols: (+)- Catechin hydrate ≥ 98% (C) (C1251), (-)- Epigallocatechin gallate 95% from green tea (EGCG) (E4143), Chalcones: Phloretin ≥ 99% (PTIN) (P7912) and Phloridzin dihydrate from apple wood, ≥99% (PDIN) (P3449), Isoflavones: Genistein ≥ 98% (GE) (G6649), Flavone ≥99.0% (FLA) (F2003), and the Hydrolysable Tannins (HTs): Gallic acid (GA) (G7384) and Tannic acid (TA) (T0200). All compounds were diluted either in Phosphate Buffer Solution (PBS) or Methanol (max 0,002% final concentration) depending on their solubility.

### Nematodes and Bacterial source maintenance

*Caenorhabditis elegans, wild-type* Bristol strain, N2 line was acquired from the Institute of Molecular Genetics (IGMM), CNRS, UMR-5535, 34293 Montpellier, France. Nematodes were cultured and stocked frozen until used following the standard protocols (Brenner, 1974). Identification of worm stages and transferences were performed under the microscope LEIKA EZ4D equipped with a transmitted light source. The bacterial food resource was the uracil-requiring mutant *Escherichia coli* OP50 obtained initially from Dr. J.P Nougayrede Laboratory. Prior to each assay, *E. coli* OP50 was freshly growing in LB broth medium following the routine protocols (Stiernagle, 2006; Weaver et al., 2017) at 37^a^C.

### Nematodes Grown Medium preparation

For nematode S liquid Medium for *C. elegans*, we followed the standard procedure proposed by Stiernagle. Solid Nematode Grown Medium (NGM) was prepared as Brenner (Brenner, 1974; Stiernagle, 2006) three days before the trial. NGM agar was poured into BD Falcon Petri plates 60mm of diameter. The entire surface was homogenously covered to avoid empty places, overnight at room temperature till solidification. Purified monomer solutions were set 24h before being added to the NGM. Afterwards, monomers stock solutions (200mM) were progressive diluted into the *E. coli* OP50 food source to reach the experimental conditions: 20μM, 10μM, and 5μM of monomer concentration into *E. coli* OP50 batch (0.6 final O.D.) in LB medium. Control negative solution established per assay was a compound-free bacterial batch of *E. coli* OP50 (0.6 O.D.) in LB medium. Once ready, the test solutions were poured into the NGM previously made, homogeneously distributed to cover the entire surface let it dry, wrapped with parafilm and stored until used (4°C). A total number of 8 experimental plates were prepared for each concentration tested to run two replicates each 24 hours.

### Maintenance of *C. elegans* and adult isolation

*C. elegans* population was grown in agar solid medium with *E. coli* OP50 as a food resource at 20°C. Three days before the beginning of the trial *C. elegans’* population was synchronized following the bleaching process proposed by Stiernagle (see (Porta-de-la-Riva et al., 2012; Stiernagle, 2006)). After bleaching, petri plates were washed up with an M9 buffer solution to collect the remnant eggs, and overnight at room temperature to reach the L1 stage. The following day M9 solution was stirring to obtain a homogenous suspension of synchronized L1. 100μL of M9 suspension was added to the experimental Petri plates per duplicate (Control, 20μM, 10μM, and 5μM monomer concentration), incubated at 20°C ±2.5 days. After the incubation 12 to 15 synchronized gravid adults (bearing 6 to 9 eggs in the uterus) (Altun et al., 2012) considered as 0 days old (t=0), were transferred using a worm pick into fresh experimental NGM plates. Adults and progeny were monitored at regular time point every 24h; values were associated with 24, 48 and 72 hours old. Original adults were then transferred into fresh NGM experimental plates to avoid the overlapping of generations. For the assessment of fecundity, offspring were counted on the agar surfaces by using squared references drawn at the bottom of the Petri-plates (1cm x 1cm wide) after staining with iodine to arrest development and to dye eggs and larvae. Assays organization and the total number of adult nematodes per monomer concentration are reported in Table 1.

**Table 1.** Assays organization, commercial purified monomers, nematodes, and solvents: Q (Quercetin), R (Rutin), C ((+)- Catechin hydrate), EGCG ((-)-Epigallocatechin gallate), PTN (Phloretin), PDN (Phloridzin Dihydrate), GE (Genistein), FLA (Flavone), GA (Gallic acid), TA (Tannic acid), G1 (original population), PBS (Phosphate Buffer Solution) and CH_4_O (Methanol 0,002%).

|  | **Assay 1**<br>Flavonoids |  | **Assay 2**<br>Flavanols |  | **Assay 3**<br>Chalcones |  | **Assay 4**<br>Isoflavone<br>Flavone |  | **Assay 5**<br>Hydrolysable<br>Tannins |  |
| --- | --- | --- | --- | --- | --- | --- | --- | --- | --- | --- |
| **20µM** | Q | R | C | EGCG | PTN | PDN | GE | FLA | GA | TA |
| **10µM** | Q | R | C | EGCG | PTN | PDN | GE | FLA | GA | TA |
| **5µM** | Q | R | C | EGCG | PTN | PDN | GE | FLA | GA | TA |
| **G1** | 12 | 12 | 12 | 12 | 12 | 12 | 12 | 12 | 15 | 15 |
| **Solvent** | PBS |  | PBS |  | CH<sub>4</sub>O |  | CH<sub>4</sub>O |  | CH<sub>4</sub>O |  |

### Measurements and data analysis

*C. elegans* first generation of worms (G1) was grown in a treatment culture media till adulthood, then daily transferred to fresh plates for 72 hours. Petri plates affected by fungi were discarded from the trial. Dead G1 nematodes and internal hatching were scored, nematodes found outside the agar surface were censored. Daily and total reproductive output and egg development rate per monomer concentration were evaluated by an independently Split-plot ANOVA repeated a measures analysis, data were not transformed (Hart, 2006). The non-parametric tests Kolmogorov-Smirnov (Normality), Kendall and Friedman tests. Lastly the Non-parametric Spearman’s Rho and Partial Correlation tests’ were run considering the total brood size of *C. elegans* exposed to every specific monomer despite concentration, and the mean rate of development (data analyses were performed using SPSS version 12.0 (SPSS, Inc., Chicago IL). The interpretation of the Spearman’s and partial correlation coefficients were made pondering: 0= zero; ±0.1-0.3 = weak, ±0.4-0.6 = moderate, ±0.7-0.9 = strong, and ±1 = perfect (Akoglu, 2018). Percentage of change on larval development rate were daily scored upon the formula: ((T *100)/ C) -100), being T = the treated G1 offspring, and C= the non-treated G1 offspring. Control values per trial were set up as “0”, positive percentages of monomers (x>0) were pondered as higher than control ones, values (x<0) were associated with a reduction on egg-hatching fertility measures. Percentages were organized into 7 different levels, estimated by nclss.sturges using R software (version 3.5.1(2018-07-02)).

## Results

Due to contamination with fungi, one of the replications of PZN 20 was discarded.

### Effects on Egg-laying rate

The Split-plot Repeated Measures arises significant differences in the effect of time Within-subjects effects on the egg-laying rate (p<0.001). Contrary, the non-parametric test Friedman and Kendall was only significance for GE-FLA (p<0.001) for the effect of time (24, 48 and 78 hours). No differences were found for the effect of time and monomer concentration interaction. Post Hoc test Multiple Comparison (Games Howell), was statistically significant (p<0.05) for C20 compared to Control and C10.

### Effects on egg hatching, percentages of eggs to larval development

The Split-plot Repeated Measures displayed significant differences for the effect of time within-subjects (p<0.05) but for EGCG-C, confirmed a posteriori by the Non-parametric test Friedman and Kendall. Time and monomer concentration interaction was significant (p<0.05) for the Hydrolysable tannins assay GA and TA, being close to significant for the flavonoids Q and R. Test of Between-subjects effects were significant (p<0.05) for Q-R Assay and GE-FLA assays respectively. Total egg output and mean development after Spearman’s Rho correlations arose negative correlations in a Moderate level for EGGC-C (p<0.05) and PTN-PDN (p<0.05), strong in the case of GA-TA (p<0.01). A positive Spearman’s Rho correlation was found for GE-FA (p<0.01) in a moderate level (Table 2).

**Table 2.** Spearman’s rho: Total egg output and Mean development per Monomer: Q (Quercetin), R (Rutin), C ((+)-Catechin hydrate), EGCG ((-)-Epigallocatechin gallate), PTN (Phloretin), PDN (Phloridzin Dihydrate), GE (Genistein), FLA (Flavone), GA (Gallic acid) and TA (Tannic acid).

|  | **Q-R** | **EGGC-C** | **PTN-PDN** | **GE-FLA** | **GA-TA** |
| --- | --- | --- | --- | --- | --- |
| Correlation Coefficient | 0,231 | -0,468 | -0,615 | 0,516 | -0,815 |
| Sig. (1-tailed) | 0,214 | 0,046 | 0,013 | 0,029 | 0,000 |
| N | 14 | 14 | 13 | 14 | 14 |
| P-value |  | * | * | ** | ** |

\* Correlation is significant at the 0.05 level (1-tailed).
\*\* Correlation is significant at the 0.01 level (1-tailed).

Partial correlation analyses showed a Negative Moderate correlation for GA-TA and a strong negative correlation for PTN and PDN (P<0.05). EGCG-C resulted close to being statistically significant (p= 0.054) (Table 3).

**Table 3.** Partial Correlation: Total egg output and Mean development per Monomer: Q (Quercetin), R (Rutin), C ((+)-Catechin hydrate), EGCG ((-)-Epigallocatechin gallate), PTN (Phloretin), PDN (Phloridzin Dihydrate), GE (Genistein), FLA (Flavone), GA (Gallic acid) and TA (Tannic acid).

| Control Variables: All |  |  |  |  |  |  |
| --- | --- | --- | --- | --- | --- | --- |
| fertilities addition |  |  |  |  |  |  |
| Partial Correlations | Average development | **Q-R** | **EGCG-C** | **PTN-PDN** | **GE-FLA** | **GA-TA** |
| Control Variables: | Correlation | -0,144 | -0,466 | -0,728 | 0,184 | -0,556 |
| Monomer concentration | Significance (1-tailed) | 0,319 | 0,054 | 0,004 | 0,274 | 0,024 |
|  | df | 11 | 11 | 10 | 11 | 11 |

**Table 4.**
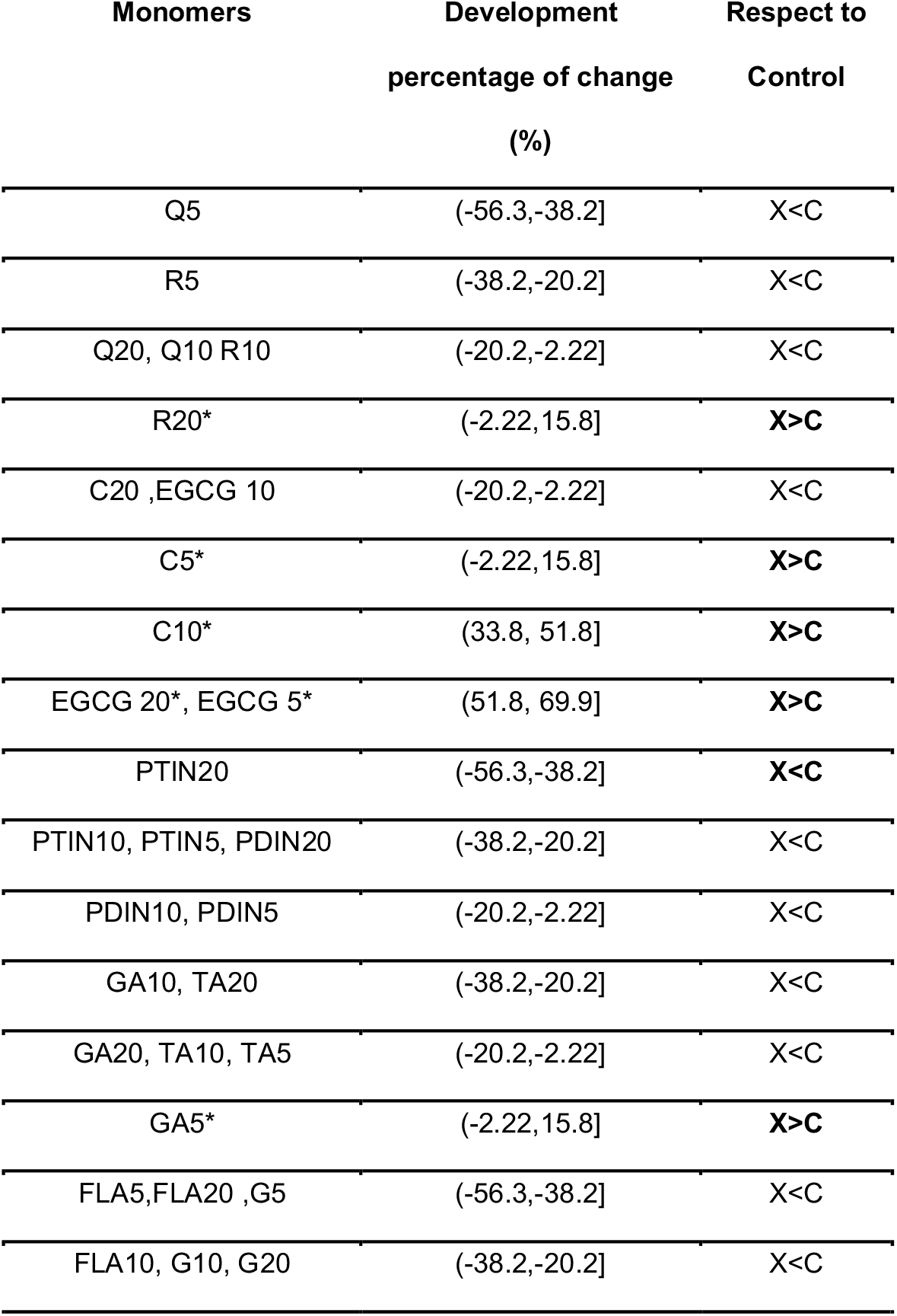
Percentages of change on egg development respect to control values. Percentages are organized into seven ranges from negative -56.3% to positive 69.9%. Signs correspond to decreases or increases on larva hatching respect to control values Q (Quercetin), R (Rutin), C ((+)-Catechin hydrate), EGCG ((-)-Epigallocatechin gallate), PTN (Phloretin), PDN (Phloridzin Dihydrate), GE (Genistein), FLA (Flavone), GA (Gallic acid) and TA (Tannic acid) and *(Higher percentages of development).

| Monomers | Development<br>percentage of change<br>(%) | Respect to<br>Control |
| --- | --- | --- |
| Q5 | (-56.3,-38.2] | X<C |
| R5 | (-38.2,-20.2] | X<C |
| Q20, Q10 R10 | (-20.2,-2.22] | X<C |
| R20* | (-2.22,15.8] | **X>C** |
| C20 ,EGCG 10 | (-20.2,-2.22] | X<C |
| C5* | (-2.22,15.8] | **X>C** |
| C10* | (33.8, 51.8] | **X>C** |
| EGCG 20*, EGCG 5* | (51.8, 69.9] | **X>C** |
| PTIN20 | (-56.3,-38.2] | **X<C** |
| PTIN10, PTIN5, PDIN20 | (-38.2,-20.2] | X<C |
| PDIN10, PDIN5 | (-20.2,-2.22] | X<C |
| GA10, TA20 | (-38.2,-20.2] | X<C |
| GA20, TA10, TA5 | (-20.2,-2.22] | X<C |
| GA5* | (-2.22,15.8] | **X>C** |
| FLA5,FLA20 ,G5 | (-56.3,-38.2] | X<C |
| FLA10, G10, G20 | (-38.2,-20.2] | X<C |

*Percentages of change on larval development* rate were organized into 7 ranges (−56.3,-38.2], (−38.2,-20.2], (−20.2,-2.22], (−2.22,15.8], (15.8,33.8], (33.8,51.8] and (51.8,69.9]. The spectrum of percentages of change compared to each blank resulted negative, but for R20, C5, C10, EGCG 20, EGCG 5, and GA5, associated with a net increase of hatched larvae (Fig 1).

**Fig 1:**
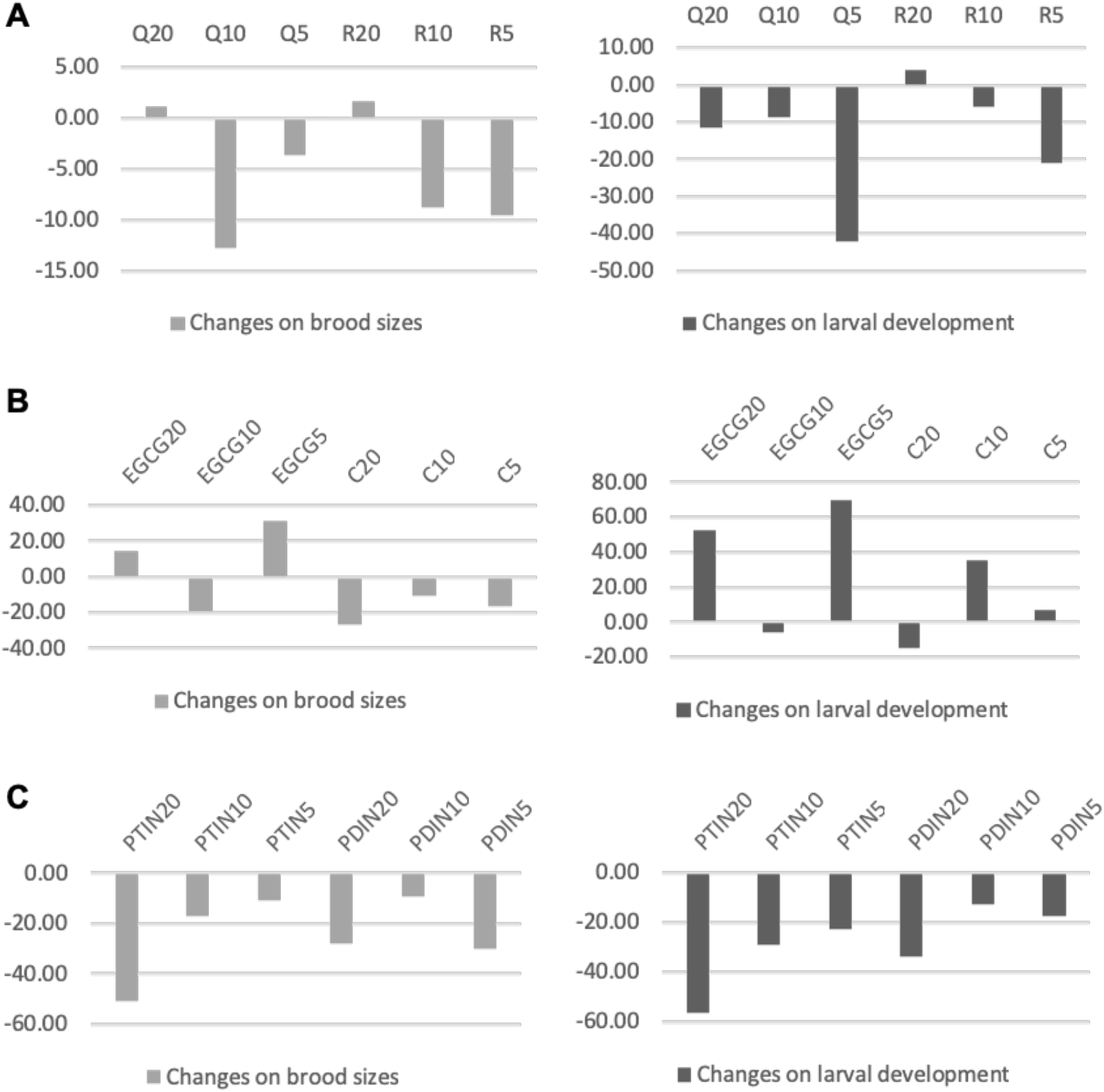
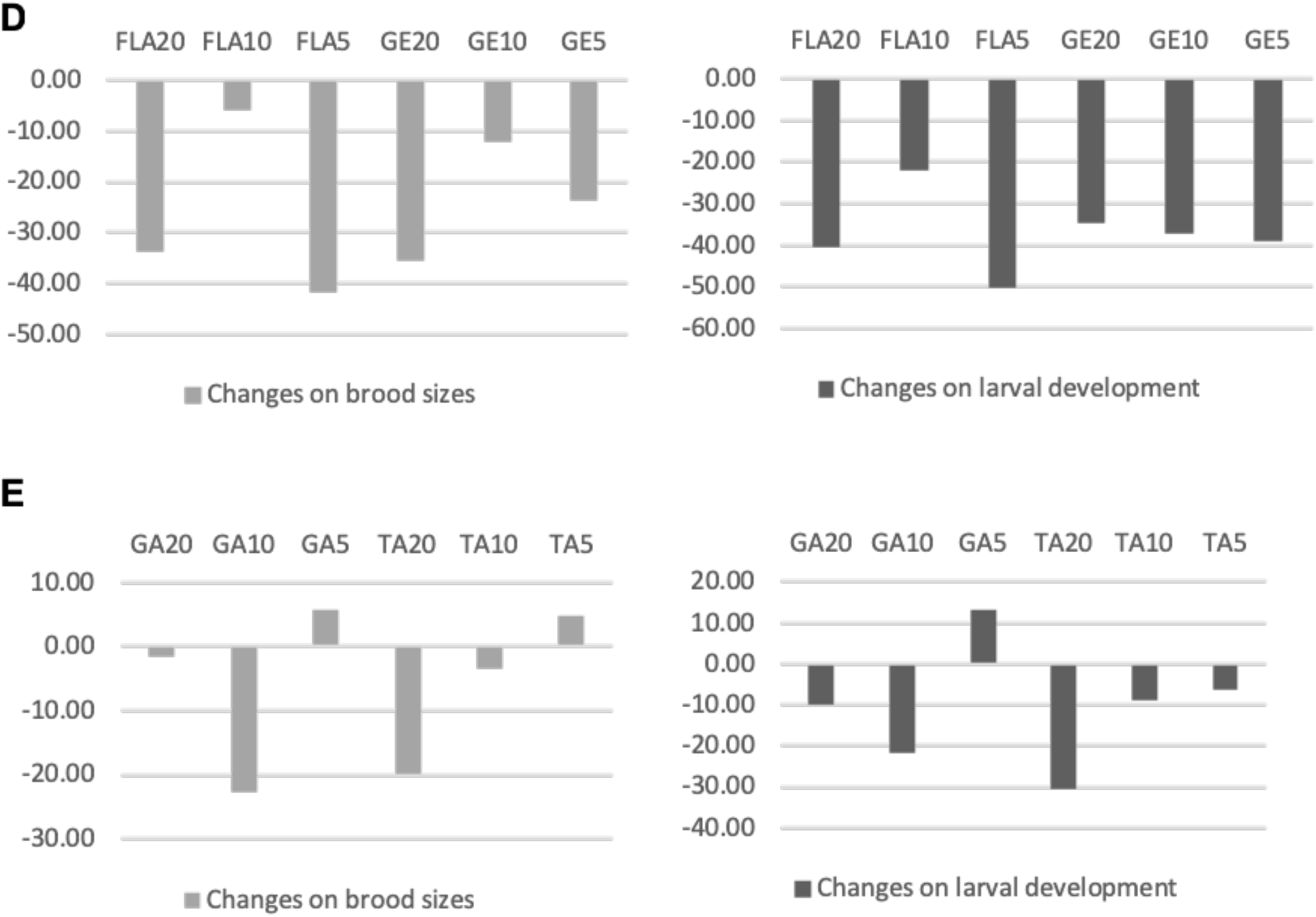
Percentages of change on *Caenorhabditis elegans* of laid eggs and eggs hatched though time. Left columns in light grey, represent percentages of change on brood sizes. Right columns in dark grey, show the percentages of change on larval development. Rows from the top to the bottom A: Q (Quercetin), R (Rutin) Assay, B: C ((+)-Catechin hydrate), EGCG ((−)- Epigallocatechin gallate) assay, C: PTN (Phloretin), PDN (Phloridzin Dihydrate) assay, D: FLA (Flavone), GE (Genistein) assay, and E: GA (Gallic acid) and TA (Tannic acid) assay.

## Discussion

The most challenging limits while screening AH effects on adult stages are: 1) maintaining alive outside the host stages adapted to live in the gastrointestinal gut, 2) reproduce *in vitro* environmental and physiological gut conditions, 3) detect anthelmintic effects and mechanism of action of compounds in adult stages *in vivo*. To set up a model to evaluate effects on the fertility of adult stages, we selected *C. elegans* model based on its multiple advantages. Literature published remarks about the benefits of liquid a medium (Katiki et al., 2011) or 3D medium (Lee et al., 2016). *C. elegans’* culture in liquid mediums is considered a low costly excellent option for a rapid proliferation in a short period. Several trials can be run simultaneously, allowing you to easily screen lethal effects on worms exposed to different monomers concentrations and solvents, evaluating potential effects on host and environment consequences of the compounds tested (Mathew et al., 2016). Thus, we first set up a protocol in a complete S Medium inoculated with *E. coli* (OD. 0.6) as a food source. Synchronized adult worms were cultured in S liquid medium in a 3×2 BD Falcon culture multiwell plates for 48h. Despite the practical conveniences, we found obstacles in the evaluation of fecundity due to technical issues to compute offspring due to the turbidity of the medium, and offspring losses after aliquoting. Eggs and larvae got stuck into the tips while micropipetting. Thus FuDR was added at the recommended dosage level to avoid lethal effects (Hope, 1999), it did not prevent from eggs and larvae went missing altering the real numbers of fertility values (Data not shown). Furthermore, we found a phenomenon of cross-contaminations of fungi between the wells of the 3×2 multiwell plates. To avoid altered offspring numbers and cross contamination we decided then to move forward solid B agar NGM. Synchronized *C. elegans* grew as fast as expected at 20°C, and worms were easily transferred to fresh plates using the worm pick, egg and larvae measurement resulted easier and more rigorous. Solid agar medium for *C. elegans* smoothed worms’ transferences decreasing potential offspring loses and contamination, but required us a minimum of 2 weeks to complete a trial. However, agar NGM plates, require longer time periods for the setting up: a minimum of 3 days to get ready NGM plates for two monomers at 3 different concentrations plus control plates, 72 hours of G1 culture, 3 additional days to run fertility test per se, and about 1 to 2 days to compute offspring. For the setting up of the protocol for fertility measures, selected polyphenols were diluted into two different solvents depending on their solubility: Phosphate Buffer Solution, and methanol in less than 0,002% final concentration. Likewise, an active bacterial food resource was a key factor to maintain for *C. elegans* life cycle with no accessory variables potentially affecting life span or biological activities. Accordingly, we performed a bacterial growing test on agar squared plates (20°C) for 72h to discard any potential bactericide effects following Dr. J.P Nougayrede Laboratory protocols. No antibacterial activity was observed (data not shown). *E. coli* OP50 was then grown at the experimental monomer and solvent concentration. Either bacterial food source and G1 *C. elegans* tolerated methanol doses (Katiki et al., 2013; Katiki et al., 2011). Neither lethality nor bagging effects were found on G1 *C. elegans*. Many of the published researches on *C. elegans* found are focused on lifespan, anti-aging and antioxidant effects of polyphenolic compounds. As far as I know not too many trials have been oriented to screen the AH effects of purified monomers rather than plant extracts. Our trials indicated a great variability of results depending on each particular polyphenol and concentration tested. Contrary to published studies on the AH effects of CT and HT plant rich-extracts against *C. elegans* (Katiki et al., 2013; Spiegler et al., 2016), purified monomers did not disclosed neither lethality nor “bagging effects” (hatching inside adult hermaphrodite worms) (Bull et al., 2007; Saul et al., 2010) on G1. However, some adults were missed from the agar surface not due to deceases. We assumed potential accidentally falls during the storage or potential active evasive behaviors and discarded those adults from the trial. The analyses on brood sizes were performed then, pondering the exact number of adults transferred every 24 hours. The daily amount of eggs and larvae and the final brood size varied significantly depending on the fertility schedules (Muschiol et al., 2009). Highest offspring sizes were observed at t=48 corresponding to 108 hours of age approximately. However, we also analyzed this trend with a non-parametric test only observed significance for the flavone and the isoflavone. Observing the egg-laying behavior, G1 exposed to Flavone and Genistein showed upper output at t=72. However, at the beginning of genistein-flavones assay *C. elegans* were still young adults (not carrying eggs inside) (Sulston and Hodgkin, 1988) so we might suspect a potential shifting of the maximum output peak. All the worms tested in each independent trial came from the same original N2 strain, control tests were run in parallel to treated worms so considering our results we cannot assume direct effects of monomers on the age-specific fecundity or sexual maturity (Muschiol et al., 2009; Riddle et al., 1997). We also analyzed the potential effects of the interaction of the variable time and monomer, to discern possible effects in egg excretion but we did not find any correlation among the assay. So, according to *C. elegans* phenology and senescence at 20°C (Hart, 2006), the effect of time might be associated with the regular process of aging rather than an effect of the treatment per se. To discard aging effects on the viability of the eggs, additionally, we evaluated the effect of time within subjects. Time also resulted significant on the mean development hatching, but contrary it was not an effect within flavanols EGCG-C, so a potential effect of this monomers might be expected. Finally, we obtained a significant effect for Time and monomer concentration interaction for the Hydrolysable tannins assay GA and TA, close to significance for the flavonoids Q and R. The evaluation of effects on the reproduction of *C. elegans* as a potential tool to extrapolate to parasitic nematodes species in order to decrease reinfections and the contamination of the pastures. Considering that *C. elegans* parthenogenetic females, self-fertilized their ova, effects on reproductions can be associated with egg production, inhibitory effects in egg-laying, hatching, development or post-hatching effects on the offspring (Bull et al., 2007). Our first trial screened potential AH effects of the flavonoids Quercetin and Rutin. Quercetin was associated with in vitro improvements of moxidectin bioavailability in blood samples from lambs (Dupuy et al., 2003). Similarly, extracts of Sainfoin rich in 3-flavonol glycosides as Rutin inhibited L3 larval migration of *H. contortus* in vitro (Barrau et al., 2005). Nevertheless, Quercetin did not show any effect on the reproductive parameters on the nematode *C. elegans* (Pietsch et al., 2009) but enhanced its lifespan and resistance to oxidative stress (Kampkötter et al., 2008; Pietsch et al., 2009). Within our trial, we notice flavonoids potentially decrease brood sizes and percentage of development, except for 20μM concentration. Quercetin and Rutin 20μM seems to stimulate reproduction in terms of egg production, or both parameters respectively. During our second assay, we evaluated the Flavanols Catechin and Epigallocatechin gallate, both precursors of condensed tannins. Catechin is a procyanidin (Mueller-Harvey, 2006) while Epigallocatechin-gallate is a galloyl-derivate (Hagerman, 2002). Studies on the anthelmintic effects of CTs against GINs showed a dependency on the structure and AH activity (Aerts et al., 1999; Mueller-Harvey, 2006). Procyanidin monomers have less AH effectiveness than prodelphinidins or galloyl-derivatives (Brunet and Hoste, 2006; Brunet et al., 2008a). Contrary, our studies showed better outcomes for procyanidin rather than the galloyl-derivate. Catechin revealed a general decrease of brood size, significant for 20μM concentration, and a decreased of 15% on egg hatching respect to control values. This also contrasts with previous studies on *C. elegans* not confirming any effect of Catechin on reproduction (Saul et al., 2009; Surco-Laos et al., 2012). In the case of the epigallocatechin-gallate, brood sizes and egg-hatching of *C. elegans* seems to be stimulated. EGCG10μM decreased reproduction but with no in a significative manner. The general pattern for this monomer, seems to be associated with a net increase of fertility and percentages of hatching (EGCG20 (53%), EGCG 5 (70%)). Our results seems to be in line with recent publication on *C. elegans* with no inhibition of egg hatching or larval migration for B-type proanthocyanidins rich extract from *Paullinia pinnata* roots (Spiegler et al., 2016) or the lifespan and anti-aging effects of EGCG in *C. elegans* (Brown et al., 2006; Zhang et al., 2009). The effects of the chalcones Phloretin and Phloridzin, a Glycosylated phloretin (Gosch, 2010; Gosch et al.) were also screened. Previous studies remarked the bioactivity of dihydrochalcones as functional antioxidants (Dugé de Bernonville et al., 2010). We discarded one of the replications of PZN 20; despite this, we detected a general decrease in the fertility of G1 (egg output and percentages of development). Results on Genistein and flavone indicated a significant general decrease in reproductive parameters. Egg output and development were positively correlated in a moderate level. The nematocidal effects of Genistein against helminth parasite and cestodes were already confirmed (Tandon and Das, 2018), as well as the embryonic and larval lethality of Flavone against *C. elegans* (Yong-Uk et al., 2008). Due to the low number of studies on Hydrolysable tannin, we decided to include Gallic and Tannic acid among our assays. We found a strong negative correlation between egg-hatching and egg-laying for both monomers; the reason why an increase in the egg-hatching might compensate for the relative decrease in egg-laying behavior. Tannic acid indicated effectiveness against egg-hatching. The spectrum of percentages of change compared to each blank resulted negative, but for GA5, associated with a net increase of larvae hatched. This contrast with other studies where TA in a low concentration did not modulate daily reproduction patterns and total brood size of *C. elegans* (Saul et al., 2010).

In summary, we found solid agar Nematode Growth medium as the appropriate medium to screen changes in fecundity of *C. elegans* once the anthelmintic is added to the food resource. We were able to evaluate the main effects of purified monomers on functional activities as egg-laying behavior and egg-hatching in an easier and reduced costs.. Due to the low number of replications. we found difficulties in data analyses and interpretation. To overcome this issue, we suggest increasing the number of replications per treatment, to get a more powerful statistical analysis. However, this will also require a better and higher level of organization. View our results we suggest to deeper analyze the effects of chalcones phloretin, Phloridzin, as well as flavone and genistein as potential AHs in the modulation of fecundity. In terms of flavonols, effectiveness seems to follow a dose-dependent patterns. Effects of time schedule and monomer concentration were found for the Hydrolysable tannins Gallic and Tannic acid in reproductive parameters. Tannic acid showed effectiveness against egg-hatching. Likewise, we propose inquiring about potential accumulative effects on the offspring in a long-term analyzing F2 and F3 generations.

## Conclusions

The solid Nematode Growth Medium offered us a proper reproducibly culture milieu, in considerable low time and cost manner, to screen AH effects on fertility (egg-laying and egg-hatching) behavior. We found a nematocidal trend on the reproductive parameters of *C. elegans* treated with Phloretin, Phloridzin, and flavone. In a dose-dependent manner, monomers as Catechin might be helpful in the modulation of fertility. A minimum of three replications per monomer concentration is suggested to get a more robust statistic result.

## Acknowledgments and additional information

This work was partially supported by the author’s personal private funds and Pôle Emploi benefits for the unemployed. The present manuscript was included as a chapter of the corresponding author Ph.D. manuscript submitted in 2015. I would like to thank Dr. Hervé Hoste to allow me to work at his laboratory at the ENVT, to Dr. J.P Nougayrede to generously allow me to use his laboratory for the microbiological cultures. A special mention to Ms. Michèle Boury for her valuable help in the establishment of the protocols and technical support. Dr. Daniel Paredes from the University of Coimbra for his comments and suggestions. The mention of commercial products is merely informative; no funds were received from them. The author, claims for the improvement of labor conditions at research.

## Notes

**Conflict of interest:** Author declares no conflict of interest.

### Competing Interest Statement

I was working with no funds during part of this trial and PhD. Scarce guidance was received. Part of the experiments were performed by me while I was irregularly working, surviving, thanks to unemployment governmental funds and my own savings. This manuscript is included within my Ph.D. manuscript, available at Universidad Autonoma de Madrid online repository, since 2015.

## References

Aerts, R.J., Barry, T.N., McNabb, W.C., 1999. Polyphenols and agriculture: beneficial effects of proanthocyanidins in forages. Agriculture, Ecosystems and Environment 75, 1–12.

Akoglu, H., 2018. User’s guide to correlation coefficients. Turkish journal of emergency medicine 18, 91–93.

Altun, Z.F., Herndon, L.A., Wolkow, C.A., Crocker, C., Lints, R., Hall, D.H. 2002-2019. Wormatlas. http://www.wormatlas.org.

Arroyo-Lopez, C., Manolaraki, F., Saratsis, A., Saratsi, K., Stefanakis, A., Skampardonis, V., Voutzourakis, N., Hoste, H., Sotiraki, S., 2014. Anthelmintic effect of carob pods and sainfoin hay when fed to lambs after experimental trickle infections with Haemonchus contortus and Trichostrongylus colubriformis. Parasite 21, 71–71.

Athanasiadou, S., Kyriazakis, I., Giannenas, I., Papachristou, T.G., 2009. Nutritional consequences on the outcome of parasitic challenges on small ruminants, Méditerranéennes: Série A. Séminaires Méditerranéens 85, 29–40.

Barrau, E., Fabre, N., Fouraste, I., Hoste, H., 2005. Effect of bioactive compounds from sainfoin (Onobrychis viciifolia Scop.) on the in vitro larval migration of Haemonchus contortus: role of tannins and flavonol glycosides. Parasitology 131, 531–538.

Beynon, S.A., 2012. Potential environmental consequences of administration of anthelmintics to sheep. Veterinary Parasitology 189, 113–124.

Brenner, S., 1974. The genetics of Caenorhabditis elegans. Genetics 77, 71–94.

Brown, M.K., Evans, J.L., Luo, Y., 2006. Beneficial effects of natural antioxidants EGCG and [alpha]-lipoic acid on life span and age-dependent behavioral declines in Caenorhabditis elegans. Pharmacology Biochemistry and Behavior 85, 620–628.

Brunet, S., Hoste, H., 2006. Monomers of condensed tannins affect the larval exsheathment of parasitic nematodes of ruminants. Journal of Agricultural and Food Chemistry 54, 7481–7487.

Brunet, S., Jackson, F., Hoste, H., 2008a. Effects of sainfoin (Onobrychis viciifolia) extract and monomers of condensed tannins on the association of abomasal nematode larvae with fundic explants. International Journal for Parasitology 38, 783–790.

Brunet, S., Martinez-Ortiz de Montellano, C., Torres-Acosta, J.F.J., Sandoval-Castro, C.A., Aguilar-Caballero, A.J., Capetillo-Leal, C., Hoste, H., 2008b. Effect of the consumption of Lysiloma latisiliquum on the larval establishment of gastrointestinal nematodes in goats. Veterinary Parasitology 157, 81–88.

Bull, K., Cook, A., Hopper, N.A., Harder, A., Holden-Dye, L., Walker, R.J., 2007. Effects of the novel anthelmintic emodepside on the locomotion, egg-laying behaviour and development of Caenorhabditis elegans. International Journal for Parasitology 37, 627–636.

Bürglin, T.R., Lobos, E., Blaxter, M.L., 1998. Caenorhabditis elegans as a model for parasitic nematodes. International Journal for Parasitology 28, 395–411.

Burns, A.R., Luciani, G.M., Musso, G., Bagg, R., Yeo, M., Zhang, Y., Rajendran, L., Glavin, J., Hunter, R., Redman, E., Stasiuk, S., Schertzberg, M., Angus McQuibban, G., Caffrey, C.R., Cutler, S.R., Tyers, M., Giaever, G., Nislow, C., Fraser, A.G., MacRae, C.A., Gilleard, J., Roy, P.J., 2015. Caenorhabditis elegans is a useful model for anthelmintic discovery. Nature Communications 6, 7485.

Dugé de Bernonville, T., Guyot, S., Paulin, J.-P., Gaucher, M., Loufrani, L., Henrion, D., Derbré, S., Guilet, D., Richomme, P., Dat, J.F., Brisset, M.-N., 2010. Dihydrochalcones: Implication in resistance to oxidative stress and bioactivities against advanced glycation end-products and vasoconstriction. Phytochemistry 71, 443–452.

Dupuy, J., Larrieu, G., Sutra, J.F., Lespine, A., Alvinerie, M., 2003. Enhancement of moxidectin bioavailability in lamb by a natural flavonoid: quercetin. Veterinary Parasitology 112, 337–347.

Durmic, Z., Blache, D., 2012. Bioactive plants and plant products: Effects on animal function, health and welfare. Animal Feed Science and Technology 176, 150–162.

Fabricant, D.S., Farnsworth, N.R., 2001. The value of plants used in traditional medicine for drug discovery. Environmental health perspectives 109 Suppl 1, 69–75.

Fetterer, R.H., 1989. The cuticular proteins from free-living and parasitic stages of Haemonchus contortus. Isolation and partial characterization. Comparative Biochemistry and Physiology Part B: Comparative Biochemistry 94, 383–388.

Geary, T.G., Thompson, D.P., 2001. Caenorhabditis elegans: how good a model for veterinary parasites? Veterinary Parasitology 101, 371–386.

Gosch, C., 2010. Phloridzin: Biosynthesis, distribution and physiological relevance in plants. Phytochemistry 71, 838–843.

Gosch, C., Halbwirth, H., Schneider, B., HÃ¶lscher, D., Stich, K., 2010. Cloning and heterologous expression of glycosyltransferases from Malus x domestica and Pyrus communis, which convert phloretin to phloretin 2-O-glucoside (phloridzin). Plant Science 178, 299–306.

Hagerman, A.E. 2002. Tannin Handbook (Miami University, Oxford OH 45056).

Hart, A.C., 2006. Behavior, In: Wormbook. The C. elegans Research community., Massachusetts General Hospital Cancer Center, Charlestown, MA 02129 USA.

Hiepe, T., Lucius, R., Gottstein, B., 2006. Parasitología general. Con principios de inmunología, diagnóstico y lucha antiparasitaria, Editorial ACRIBIA, S.A. Edition Zaragoza (España), 600 p.

Holden-Dye, L., Walker, R.J., 2014. Anthelmintic drugs and nematicides: studies in Caenorhabditis elegans. WormBook, ed. The C. elegans Research Community, WormBook, doi/10.1895/wormbook.1.143.2, http://www.wormbook.org.

Hope, I.A., 1999. C. elegans. A practical approach Kindle Edition, 288 p.

Horvat, A.J.M., Babić, S., Pavlović, D.M., Ašperger, D., Pelko, S., Kaštelan-Macan, M., Petrović, M., Mance, A.D., 2012. Analysis, occurrence and fate of anthelmintics and their transformation products in the environment. Trends in Analytical Chemistry 31, 61–84.

Hoste, H., Jackson, F., Athanasiadou, S., Thamsborg, S.M., Hoskin, S.O., 2006. The effects of tannin-rich plants on parasitic nematodes in ruminants. Trends in Parasitology 22, 253–261.

Hoste, H., Martinez-Ortiz-De-Montellano, C., Manolaraki, F., Brunet, S., Ojeda-Robertos, N., Fourquaux, I., Torres-Acosta, J.F.J., Sandoval-Castro, C.A., 2012. Direct and indirect effects of bioactive tannin-rich tropical and temperate legumes against nematode infections. Veterinary Parasitology 186, 18–27.

Hoste, H., Torres-Acosta, J.F.J., Sandoval-Castro, C.A., Mueller-Harvey, I., Sotiraki, S., Louvandini, H., Thamsborg, S.M., Terrill, T.H., 2015. Tannin containing legumes as a model for nutraceuticals against digestive parasites in livestock. Veterinary Parasitology 212, 5–17.

Hu, Y., Ellis, B.L., Yiu, Y.Y., Miller, M.M., Urban, J.F., Shi, L.Z., Aroian, R.V., 2013. An Extensive Comparison of the effect of anthelmintic classes on diverse Nematodes. PLoS One 8, e70702.

Jackson, F., Varady, M., Bartley, D.J., 2012. Managing anthelmintic resistance in goats_Can we learn lessons from sheep? Small Ruminant Research 103, 3–9.

Kampkötter, A., Timpel, C., Zurawski, R.F., Ruhl, S., Chovolou, Y., Proksch, P., Wätjen, W., 2008. Increase of stress resistance and lifespan of Caenorhabditis elegans by quercetin. Comparative Biochemistry and Physiology Part B: Biochemistry and Molecular Biology 149, 314–323.

Kaplan, R.M., Vidyashankar, A.N., 2012. An inconvenient global truth worming and anthelmintic resistance. Veterinary Parasitology 186, 70–78.

Katiki, L. M., Ferreira, J.F.S., Gonzalez, J.M., Zajac, A.M., Lindsay, D.S., Chagas, A.C.S., Amarante, A.F.T., 2013. Anthelmintic effect of plant extracts containing condensed and hydrolyzable tannins on Caenorhabditis elegans, and their antioxidant capacity. Veterinary Parasitology 192, 218–227.

Katiki, L.M., Ferreira, J.F.S., Zajac, A.M., Masler, C., Lindsay, D.S., Chagas, A.C.S., Amarante, A.F.T., 2011. Caenorhabditis elegans as a model to screen plant extracts and compounds as natural anthelmintics for veterinary use. Veterinary Parasitology 182, 264–268.

Ketzis, J.K., Vercruysse, J., Stromberg, B.E., Larsen, M., Athanasiadou, S., Houdijk, J.G.M., 2006. Evaluation of efficacy expectations for novel and non-chemical helminth control strategies in ruminants. Veterinary Parasitology 139, 321–335.

Laing, R., Kikuchi, T., Martinelli, A., Tsai, I.J., Beech, R.N., Redman, E., Holroyd, N., Bartley, D.J., Beasley, H., Britton, C., Curran, D., Devaney, E., Gilabert, A., Hunt, M., Jackson, F., Johnston, S.L., Kryukov, I., Li, K., Morrison, A.A., Reid, A.J., Sargison, N., Saunders, G.I., Wasmuth, J.D., Wolstenholme, A., Berriman, M., Gilleard, J.S., Cotton, J.A., 2013. The genome and transcriptome of Haemonchus contortus, a key model parasite for drug and vaccine discovery. Genome biology 14, R88–R88.

Lee, T.Y., Yoon, K.-h., Lee, J.I., 2016. NGT-3D: a simple nematode cultivation system to study Caenorhabditis elegans biology in 3D. Biology Open 5, 529–534.

Manolaraki, F., Sotiraki S., Stefanakis A., Skampardonis V., Volanis M., Hoste H., 2010. Anthelmintic activity of some Mediterranean browse plants against parasitic nematodes. Parasitology 137, 685–696.

Martínez-Ortíz-de-Montellano, C., Arroyo-López, C., Fourquaux, I., Torres-Acosta, J.F.J., Sandoval-Castro, C.A., Hoste, H., 2013. Scanning electron microscopy of Haemonchus contortus exposed to tannin-rich plants under in vivo and in vitro conditions. Experimental Parasitology 133, 281–286.

Martínez-Ortíz-de-Montellano, C., Vargas-Magaña, J.J., Canul-Ku, H.L., Miranda-Soberanis, R., Capetillo-Leal, C., Sandoval-Castro, C.A., Hoste, H., Torres-Acosta, J.F.J., 2010. Effect of a tropical tannin-rich plant Lysiloma latisiliquum on adult populations of Haemonchus contortus in sheep. Veterinary Parasitology 172, 283–290.

Mathew, M.D., Mathew, N.D., Miller, A., Simpson, M., Au, V., Garland, S., Gestin, M., Edgley, M.L., Flibotte, S., Balgi, A., Chiang, J., Giaever, G., Dean, P., Tung, A., Roberge, M., Roskelley, C., Forge, T., Nislow, C., Moerman, D., 2016. Using C. elegans Forward and Reverse Genetics to Identify New Compounds with Anthelmintic Activity. PLoS neglected tropical diseases 10, e0005058.

McGaw, L.J., Van der Merwe, D., Eloff, J.N., 2007. In vitro anthelmintic, antibacterial and cytotoxic effects of extracts from plants used in South African ethnoveterinary medicine. The Veterinary Journal 173, 366–372.

Mitreva, M., Blaxter, M.L., Bird, D.M., McCarter, J.P., 2005. Comparative genomics of nematodes. Trends in Genetics 21, 573–581.

Molan, A.L., Duncan, A.J., Barry, T.N., McNabb, W.C., 2003. Effects of condensed tannins and crude sesquiterpene lactones extracted from chicory on the motility of larvae of deer lungworm and gastrointestinal nematodes. Parasitology International 52, 209–218.

Mueller-Harvey, I., 2006. Unravelling the conundrum of tannins in animal nutrition and health. Journal of the Science of Food and Agriculture 86, 2010–2037.

Muschiol, D., Schroeder, F., Traunspurger, W., 2009. Life cycle and population growth rate of Caenorhabditis elegans studied by a new method. BMC ecology 9, 14–14.

Nguyen, T.M., Binh, D.V., Ãrskov, E.R., 2005. Effect of foliages containing condensed tannins and on gastrointestinal parasites. Animal Feed Science and Technology 121, 77–87.

Pietsch, K., Saul, N., Menzel, R., Stürzenbaum, S., Steinberg, C., 2009. Quercetin mediated lifespan extension in Caenorhabditis elegans is modulated by age-1, daf-2, sek-1 and unc-43. Biogerontology 10, 565–578.

Porta-de-la-Riva, M., Fontrodona, L., Villanueva, A., Cerón, J., 2012. Basic Caenorhabditis elegans methods: synchronization and observation. Journal of visualized experiments, 4019.

Rand, J.B., Johnson, C.D., 1995. Chapter 8 Genetic Pharmacology: Interactions between Drugs and Gene Products in Caenorhabditis elegans, In: Epstein, H.F., Shakes, D.C. (Eds.) Methods in Cell Biology. Academic Press, pp. 187–204.

Riddle, D., Blumenthal, T., Meyer, B., 1997. Section III Aging in C. elegans, In: C.elegans II Cold Spring Harbor Laboratory Press, NY, p. 1222.

Rochfort, S., Parker, A.J., Dunshea, F.R., 2008. Plant bioactives for ruminant health and productivity. Phytochemistry 69, 299–322.

Sangster, N.C., Gill, J., 1999. Pharmacology of Anthelmintic Resistance. Parasitology Today 15, 141–146.

Saul, N., Pietsch, K., Menzel, R., Stürzenbaum, S.R., Steinberg, C.E.W., 2009. Catechin induced longevity in C. elegans: From key regulator genes to disposable soma. Mechanisms of Ageing and Development 130, 477–486.

Saul, N., Pietsch, K., Menzel, R., Stürzenbaum, S.R., Steinberg, C.E.W., 2010. The longevity effect of tannic acid in Caenorhabditis elegans: Disposable Soma meets hormesis. The Journals of Gerontology: Series A 65A, 626–635.

Smith, R., Pontiggia, L., Waterman, C., Lichtenwalner, M., Wasserman, J., 2009. Comparison of motility, recovery, and methyl-thiazolyl-tetrazolium reduction assays for use in screening plant products for anthelmintic activity. Parasitology Research 105, 1339–1343.

Spiegler, V., Liebau, E., Hensel, A., 2017. Medicinal plant extracts and plant-derived polyphenols with anthelmintic activity against intestinal nematodes. Natural Product Reports 34, 627–643.

Spiegler, V., Peppler, C., Werne, S., Heckendorn, F., Sendker, J., Liebau, E., Agyare, C., Hensel, A., 2016. Anthelmintic activity of a traditionally used root extract from Paullinia pinnata. Planta Medica 81, S1–S381.

Stiernagle, T. 2006. Worm Book. The oline review of C.elegans Biology. In Maintenance of C. elegans (Caenorhabditis Genetics Center, University of Minnesota, Minneapolis, MN 55455 USA).

Strange, K., 2006. C. elegans: Methods and Applications, Vol 351. Human Press Inc., Totowa, NJ.

Sulston, J., Hodgkin, J., 1988. The Nematode Caenorhabditis elegans. W.B. Wood, ed. Cold Spring Harbor Laboratory Press, New York, 587 p.

Surco-Laos, F., Dueñas, M., González-Manzano, S., Cabello, J., Santos-Buelga, C., González-Paramás, A.M., 2012. Influence of catechins and their methylated metabolites on lifespan and resistance to oxidative and thermal stress of Caenorhabditis elegans and epicatechin uptake. Food Research International 46, 514–521.

Tandon, V., Das, B., 2018. Genistein: is the multifarious botanical a natural anthelmintic too? Journal of Parasitic Diseases 42, 151–161.

Tang, S.Y., Halliwell, B., 2010. Medicinal plants and antioxidants: What do we learn from cell culture and Caenorhabditis elegans studies? Biochemical and Biophysical Research Communications 394 1–5.

Thompson, D.P., Klein, R.D., Geary, T.G., 1996. Prospects for rational approaches to anthelmintic discovery. Parasitology 113, 217–238.

Torres-Acosta, J.F.J., Hoste, H., 2008. Alternative or improved methods to limit gastro-intestinal parasitism in grazing sheep and goats. Small Ruminant Research 77, 159–173.

Vargas-Magaña, J.J., Torres-Acosta, J.F.J., Aguilar-Caballero, A.J., Sandoval-Castro, C.A., Hoste, H., Chan-Pérez, J.I., 2014. Anthelmintic activity of acetone–water extracts against Haemonchus contortus eggs: Interactions between tannins and other plant secondary compounds. Veterinary Parasitology 206, 322–327.

Wagil, M., Białk-Bielińska, A., Puckowski, A., Wychodnik, K., Maszkowska, J., Mulkiewicz, E., Kumirska, J., Stepnowski, P., Stolte, S., 2015. Toxicity of anthelmintic drugs (fenbendazole and flubendazole) to aquatic organisms. Environmental science and pollution research international 22, 2566–2573.

Waller, P.J. 2007. Antihelmintics and resistances: A review (Novartis Animal Health Inc. Basel, Switzerland).

Waterman, C., Smith, R.A., Pontiggia, L., DerMarderosian, A., Anthelmintic screening of Sub-Saharan African plants used in traditional medicine. Journal of Ethnopharmacology 127, 755–759.

Weaver, K.J., May, C.J., Ellis, B.L., 2017. Using a health-rating system to evaluate the usefulness of Caenorhabditis elegans as a model for anthelmintic study. PLoS One 12, e0179376.

Whelan, M., Kennedy, D.G., Trigueros, G., Cannavan, A., Boon, P.E., Wapperom, D., Danaher, M., 2012. Anthelmintic drug residues in beef: UPLC-MS/MS method validation, European retail beef survey, and associated exposure and risk assessments AU - Cooper, K.M. Food Additives & Contaminants: Part A 29, 746–760.

Wood, W.B., 1988. The Nematode Caenorhabditis elegans, Vol 17 University of Colorado, Boulder, 667 p.

Yong-Uk, L., Ichiro, K., Yoongho, L., Wan-Suk, O., Young-Ki, P., and Yhong-Hee, S., 2008. Inhibition of Developmental Processes by Flavone in Caenorhabditis elegans and Its Application to the Pinewood Nematode, Bursaphelenchus xylophilus. Molecules and Cells 26, 171–174.

Zhang, L., Jie, G., Zhang, J., Zhao, B., 2009. Significant longevity-extending effects of EGCG on Caenorhabditis elegans under stress. Free Radical Biology and Medicine 46, 414–421.

